# CRESSENT: a Bioinformatic Toolkit to Explore and Improve ssDNA Virus Annotation

**DOI:** 10.1101/2025.07.14.664782

**Authors:** R.R. Pavan, M.B. Sullivan, M.J. Tisza

## Abstract

Single-stranded DNA (ssDNA) viruses are important components of diverse ecosystems, however, it remains challenging to systematically identify and classify them. This is in part due to their broad host range and resulting genomic diversity, structure, and rapid evolution rates. In addition, distinguishing genuine ssDNA genomes from contaminating sequences in metagenomic datasets (e.g., from commercial kits) has been an unresolved issue for years. Here, we present **CRESSENT** (**CRESS**-DNA **E**xtended a**N**notation **T**oolkit), a comprehensive and modular bioinformatic pipeline focused on ssDNA virus genome-to-analysis and annotation. The pipeline integrates multiple functionalities organized into six modules: sequence dereplication, decontamination, phylogenetic analysis, motif discovery, stem-loop structure prediction, and recombination detection. Each module can be used independently or in combination with others, allowing researchers to customize their analysis workflow. With this tool, researchers can comprehensively and systematically include ssDNA viruses in their viromics workflows and facilitate comparative genomic studies, which are often limited to dsDNA viruses, therefore leaving behind a crucial component of the microbiome community under study.

## 1. BACKGROUND

Single-stranded DNA (ssDNA) viruses are among the most abundant and diverse biological entities on Earth, yet they remain one of the least understood groups of viruses. These small-genome viruses infect hosts across all domains of life, archaea, bacteria, and eukaryotes, and have been found in nearly every environment examined, from human gut, terrestrial soils and oceans to extreme ecosystems like hydrothermal vents and Antarctic lakes (Bezuidt and Makhalanyane, 2024; Kazlauskas et al., 2019; Kim et al., 2011; Rosario et al., 2018). Their ecological roles are broad: they influence host population dynamics, drive genome evolution through horizontal gene transfer, and modulate microbial community structure (Kazlauskas et al., 2019; Qazi, 2016; Zhao et al., 2019). In clinical and agricultural contexts, ssDNA viruses are highly consequential. Pathogens like Tomato yellow leaf curl virus and Banana bunchy top virus cause billions in crop losses annually (Golyaev et al., 2025; Qazi, 2016), while in humans, ssDNA phages such as Microviridae members shape gut microbiota composition with implications for inflammation and disease (Creasy et al., 2018).

The advent of metagenomic sequencing has led to a dramatic increase in the discovery of novel ssDNA viruses in diverse environments (Krupovic et al., 2020; Tisza et al., 2021, 2020). Despite their importance, analysis of ssDNA virus sequences remains difficult without in-depth ssDNA genomics expertise, which can vary significantly between host groups (Kazlauskas et al., 2019). Analytical challenges include their small genome size (which can often lead to sequence removal in viromics workflows), rapid evolution rates, frequent recombination events, and the presence of contaminating sequences in metagenomic datasets (Roux et al., 2019; Trubl et al., 2019). Furthermore, the absence of universally conserved genes complicates taxonomic classification and the comparative genomics of ssDNA viruses (Kazlauskas et al., 2018; Krupovic et al., 2015). Due to these constraints, most studies examine a set of common genes within certain ssDNA groups, including Rep and Cap genes with conserved functions and structures (Delwart and Li, 2012; Desingu and Nagarajan, 2022; Krupovic et al., 2020). Yet, these genes can be extremely divergent at the sequence level, and recombination frequently occurs between the Rep and Cap genes and, in some cases, within the Rep gene itself. Therefore, identification of these events involves phylogenetic analysis of the Cap gene, the Rep gene, and/or the nuclease and helicase domains of the Rep gene (Kazlauskas et al., 2019, Kazlauskas et al., 2017; Rosario et al., 2015). Followed by tanglegrams to visualize recombination (de la Higuera et al., 2020; Kazlauskas et al., 2017). Short motifs (e.g., Walker A and B) within the nuclease and helicase domains vary by clade, and these are often visualized as sequence logos (Kazlauskas et al., 2017). Non-coding stem-loops and iterons are additional conserved structures essential for the replication of these viruses, which can also be used to further delineate ssDNA lineages (Dai et al., 2024; de la Higuera et al., 2020; Torralba et al., 2024).

To address these challenges, we have developed CRESSENT, a modular and comprehensive bioinformatic pipeline specifically designed to analyze and annotate ssDNA virus sequences (See Fig. 1 for an overview). This tool consists of six modules, and each module can be used independently or in sequence, allowing researchers to customize their analysis workflow according to their specific research questions. Since the tool is modular, it allows for new modules to be added in the future. In this paper, we describe the functionality of each module and its integration into a cohesive analysis pipeline. With CRESSENT, researchers can enhance the efficiency and accuracy of ssDNA virus analysis and employ a standardized approach for comparative genomic studies of this important viral group.

**Fig 1:**
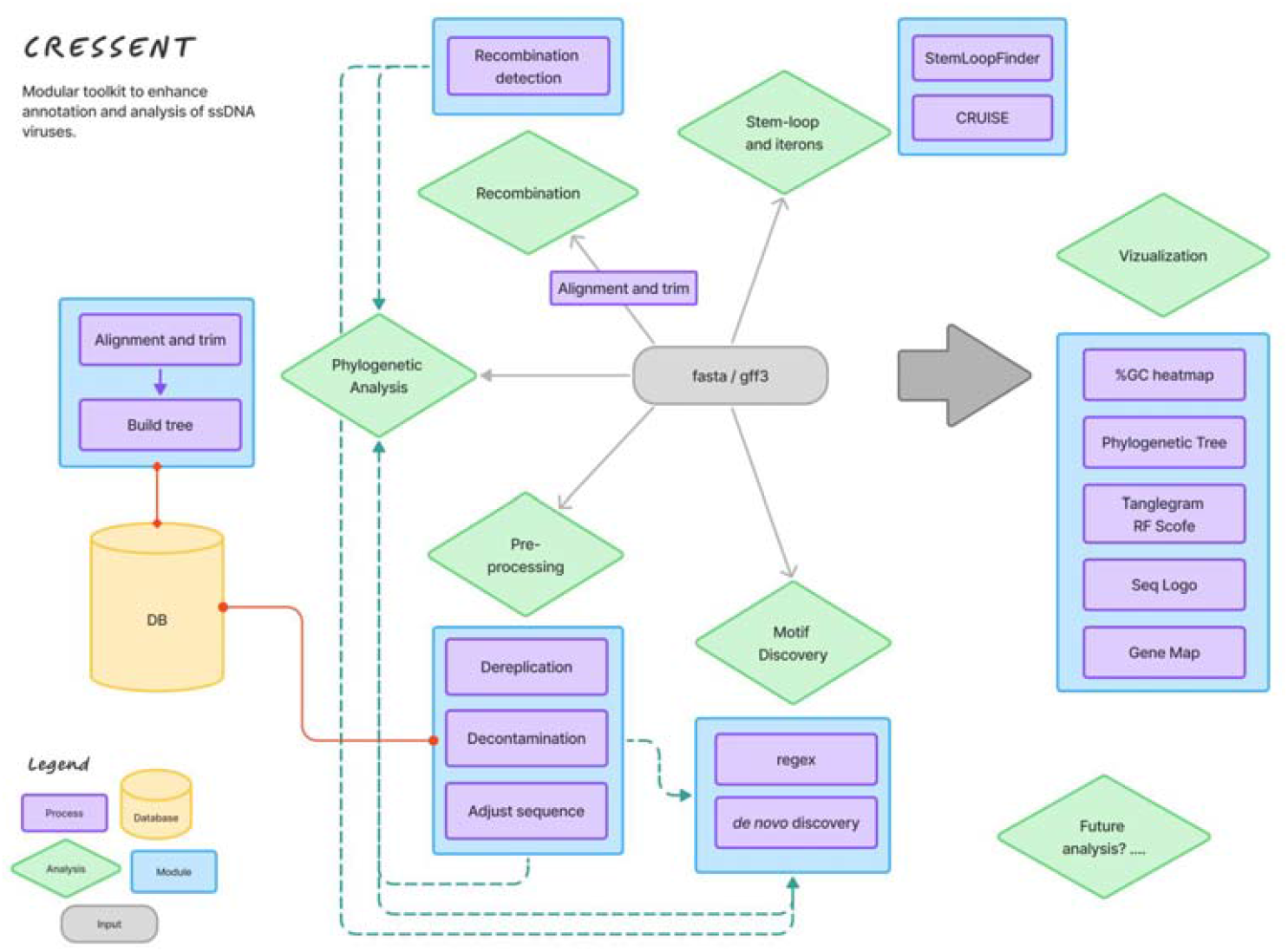
Flow chart depicting the CRESSENT modules (blue boxes), analysis (green diamonds), and processes (purple boxes). Gray and green dashed arrows indicate pipelines.

## 2. IMPLEMENTATION

The primary function of CRESSENT is to serve as an auxiliary tool to refine the annotation and detection of putative ssDNA virus sequences identified by tools such as GeNomad (Camargo et al., 2023), VirSorter2 (Guo et al., 2021), and Cenote-Taker 3 (Tisza et al., 2021). Furthermore, the integration of CRESSENT into iVirus (Bolduc et al., 2021) will facilitate the combination of different bioinformatics tools to leverage virus identification and annotation processing. The tool is written mostly in Python and R and can be used through the command line interface. The tool can be found in the GitHub repository: https://github.com/ricrocha82/cressent

The modular design of the CRESSENT allows for the flexible integration of the various components depending on the specific research questions and dataset characteristics (Fig. 1). Using the output of the virus discovery tools as input (i.e., fasta and gff files), a typical workflow might begin with dereplication to reduce dataset complexity, followed by decontamination to remove potential laboratory contaminants. The cleaned and non-redundant sequences can then be subjected to recombination detection, followed by phylogenetic analysis to understand evolutionary relationships, possibly after adjusting sequences, to begin with conserved nonanucleotide sequences using the adjust_seq.py utility. Motif discovery and annotation can provide insights into functional elements, while stem-loop and iteron annotation can identify replication-related structures. Below, we go into more detail for each of the six modules.

### 2.1 Dereplication

The dereplication module (Fig. 1) is designed to reduce sequence redundancy in datasets by clustering highly similar sequences. Sequence dereplication is widely used in viral metagenomics to manage dataset complexity and avoid bias from overrepresented sequences (Roux et al., 2019). The tool follows the current community standard for dereplication of viral contigs, as outlined by the MIUViG guidelines: 95% Average Nucleotide Identity (ANI) over at least 85% of the contig length (AF) (Roux et al., 2019). It collapses the sequences into species-level ssDNA viral operational taxonomic units (vOTUs). This module utilizes a combination of tools, including anicalc.py and aniclust.py from CheckV (Nayfach et al., 2021) and BLAST (Altschul et al., 1990), to calculate pairwise ANI and cluster sequences based on user-defined similarity thresholds. It outputs a vOTU catalog with a designated cluster representative, the full set of clustered sequences, pairwise ANI values, and supporting BLAST alignments. By removing redundancy while preserving true biological diversity, the module markedly reduces computational load for all downstream analyses.

### 2.2 Decontamination

The decontamination module (Fig. 1) addresses a critical issue in metagenomic studies: the presence of laboratory contaminants or “kitome” sequences (Olomu et al., 2020). Laboratory contamination is a concern in microbiome metagenomic studies (Duan et al., 2024; Keeler et al., 2021; Olomu et al., 2020) and can lead to the misidentification of microbial taxa, inflate diversity estimates, or result in erroneous biological interpretations if not properly controlled for through rigorous contamination-aware protocols. The systematic identification and removal of contaminant sequences is therefore essential for accurate analysis and interpretation of results.

It has been demonstrated that commercial reagents and extraction kits can contain viral DNA that may be misinterpreted as novel findings (Asplund et al. 2019). To mitigate this, a comprehensive database with 510 potential viral contaminants in laboratory reagents has been compiled (Porter et al., 2021). Contaminants included four small circular virus-like genomes (Duan et al., 2024). CRESS-like viruses were also found in laboratory reagents and may be constituents of the “kitome”. For instance, the presence of CRESS-like viruses such as Parvovirus-like Hybrid Virus (Naccache et al., 2013) and Rengasvirus (Keeler et al., 2021) in negative control samples can be mistaken for novel or biologically relevant viruses, which highlights the need to cross-reference against known reagent-associated sequences to avoid erroneous conclusions in virome and pathogen discovery research.

To limit contamination, this module uses BLAST to compare input sequences against a curated database of known contaminants derived from five published studies (Naccache et al. (2013), Asplund et al. (2019), Porter et al. (2021), Keeler et al. (2021), and Duan et al. (2024)). The module produces decontaminated sequences, decontamination statistics, and BLAST results for tracking potential contaminants. The user has full flexibility to define how stringent the decontamination process should be. To balance the removal of contaminants while retaining genuine sequences, users can fine-tune the BLAST parameters, specifically the e-value (default: 1e-10), percent identity (default: 90), and alignment coverage (default: 50).

### 2.3 Phylogenetic Analysis

The phylogenetic analysis module (Fig. 1) consists of three main components: sequence alignment, phylogenetic tree construction, and visualization/annotation of trees. The alignment component uses MAFFT (Katoh and Standley, 2013) for multiple sequence alignment and trimAl (Capella-Gutiérrez et al., 2009) for alignment trimming, while the tree construction component utilizes IQ-TREE2 for maximum likelihood phylogenetic inference. MAFFT and trimAl have been used for viral sequence alignment due to their accuracy and efficiency, which is particularly important for divergent viral sequences (Kazlauskas et al., 2018; de la Higuera et al., 2020).

Our implementation offers user-friendly customization options, including the ability to select sequences from specific viral families (e.g., Adamaviridae, Bidnaviridae, Geminiviridae) for reference and adjust alignment parameters. The module enables users to incorporate reference sequences from a built-in database of 30 ICTV-recognized ssDNA viral families (MSL39.v4), focusing on replication-associated (Rep) and capsid (Cap) proteins, or to utilize a custom reference database.

IQ-TREE2 represents the state-of-the-art in maximum likelihood phylogenetic inference and has been used extensively in viral phylogenomics (Minh et al., 2020). Similar approaches have been used to investigate the evolutionary history of CRESS-DNA viruses, as well as to examine the diversity and evolution of single-stranded DNA viruses in avian hosts (Kazlauskas et al., 2019; Chrzastek et al., 2021, Olivo et al., 2024).

Users can visualize phylogenetic trees through two dedicated modules: one for tree visualization and another for tanglegram plotting. These modules leverage specialized R packages including ggtree (Xu et al., 2022), ape (Paradis and Schliep, 2019), and dendextend (Galili, 2015). Additionally, when the tanglegram module is activated, the Robinson-Foulds (RF) distance is automatically calculated to quantify the topological dissimilarity between two phylogenetic trees. This metric provides a standardized way to assess how similar or different two trees are, which is essential for comparing phylogenetic reconstructions and evaluating the impact of different analytical methods or datasets (Briand et al., 2020).

### 2.4 MOTIF Analysis

Motif analysis is crucial for identifying functional elements in viral genomes, such as replication origins (ori), protein binding sites, and conserved protein domains. In ssDNA viruses, conserved motifs within replication-associated proteins are especially informative, as they often reflect evolutionary constraints and mechanistic roles in rolling-circle replication. For example, motifs such as the HUH endonuclease domain and the superfamily 3 (SF3) helicase domain are required for site-specific DNA cleavage and unwinding of the DNA strand, respectively. These functional domains are highly conserved across diverse ssDNA viruses, indicating strong purifying selection. As a result, their conservation across viral families can provide clues to the biochemical mechanisms of replication, robust markers for evolutionary comparisons, and taxonomic classification (Kazlauskas et al., 2018; Krupovic et al., 2020; Rosario et al., 2018, Rosario et al., 2012; Varsani et al., 2024). MEME and ScanProsite have been used for *de novo* motif discovery in viral genomics (Bailey et al., 2015; de Castro et al., 2006; Gattiker et al., 2002). For example, Tools such as MEME have been employed to explore conserved motifs in CRESS-DNA viruses, offering insights into their evolutionary relationships and classification. (Requião et al., 2020; Wang et al., 2013). Similarly, motif identification platforms like ScanProsite have been applied to viral metagenomic sequences from avian hosts to detect functionally relevant sequence patterns (Vibin et al., 2020). The pattern-based searching functionality in our tool is particularly useful for identifying known functional motifs, such as the Walker A and Walker B motifs in Rep proteins, which are indicators of rolling-circle replication mechanisms common in ssDNA viruses (Kazlauskas et al., 2018; Varsani et al., 2024; Zhao et al., 2019).

The MOTIF module (Fig. 1) enables both pattern-based motif searching and de novo motif discovery in ssDNA viral sequences. For pattern-based searching, the module uses regex and seqkit (Shen et al., 2016) to identify user-defined patterns and optionally split sequences at motif occurrences. For *de novo* motif discovery, the module integrates MEME for motif identification and optionally ScanProsite for scanning against known protein motifs. The module produces several outputs, including motif positions, sequence logos of reference and user input motifs, and genome maps to visualize the distribution of identified motifs (Hackl et al., 2024; Wagih, 2017).

### 2.5 Putative Stem Loop and Iterons Annotation

This module identifies and annotates critical noncoding secondary structures in ssDNA viral genomes: stem-loops and iterons (Fig. 1). These structures typically form the origin of replication and serve as recognition sites for viral Rep proteins (Dai et al., 2024; de la Higuera et al., 2020; D’Souza and Kool, 1992). Stem-loop structures and iterons are essential for viral replication in many ssDNA viruses. The stem-loop structure in Faba bean necrotic yellows virus (FBNYV) functions as the origin of replication, with a conserved nonanucleotide sequence within the loop being essential for initiating rolling circle replication (Timchenko et al., 1999). Additionally, a set of iterative sequences (iterons) serves as specific binding sites for Rep proteins, and their spatial arrangement plays a crucial role in determining replication efficiency in Geminiviridae. These structures represent critical functional elements that are conserved across diverse ssDNA viral families despite high sequence variability in other genomic regions (Bonnamy et al., 2023). CRESSENT includes two modules for these analyses: StemLoop-Finder for identifying DNA hairpin structures and CRUISE (CRiteria-based Uncovering of Iteron SEquences) for detecting iteron repeats that serve as recognition sites for replication proteins (Jones et al., 2022; Pratt et al., 2021). StemLoop-Finder uses the ViennaRNA library for predicting secondary structures and scores potential stem-loops based on deviation from ideal stem and loop lengths. CRUISE identifies iteron sequences, which are typically found near the origin of replication in many ssDNA viruses.

### 2.6 Recombination Detection

Recombination is a major driver of genetic diversity and evolution in ssDNA viruses (Lefeuvre et al., 2009; Martin et al., 2011). The accurate detection of recombination events is therefore crucial for understanding viral evolution, taxonomy, and epidemiology. This module identifies recombination in nucleotide sequences using a suite of methods integrated within the OpenRDP (Recombination Detection Program) framework. The analysis incorporates multiple statistical and phylogenetic approaches to improve detection sensitivity and reliability (Martin et al., 2005). The RDP method examines sequence triplets for recombination signals using a recursive segmentation algorithm, while 3Seq detects recombination based on phylogenetic incongruence among three sequences. GENECONV identifies gene conversion events by scanning for unusually long identical fragments shared between sequences. MaxChi and Chimaera both apply chi-square tests to detect breakpoints by comparing observed and expected substitutions across aligned sequences, with Chimaera using a more refined partitioning strategy. Bootscan assesses phylogenetic relationships along the sequence alignment by generating bootstrapped trees for sliding windows, highlighting regions of differing ancestry. Siscan extends this approach by calculating similarity scores across windows to detect potential recombination breakpoints. The methods implemented in this tool have been widely used in studies of viral recombination, including applications to circoviruses and parvoviruses, demonstrating their effectiveness across diverse ssDNA virus groups (Varsani et al., 2009; Fu et al., 2011). The module allows users to run specific methods or all methods together and provides options for customization through a configuration file (Fig. 1). The output includes detailed information about detected recombination events, including breakpoints, parental sequences, and statistical support.

### 2.7 Visualization and Outputs

Visualization components are integrated throughout the pipeline, including tree visualization with ggtree, and sequence logo generation for motif representation. The visualization tools make use of established R packages such as ggtree, ggtreeextra, and ggseqlogo, providing publication-quality figures for downstream use. The output files from each module are designed to be compatible with the input requirements of subsequent modules, facilitating seamless integration. Also, they can be used as inputs for other tools not included in CRESSENT. For instance, the tree and metadata files produced in the phylogenetic analysis module can be imported into online visualization tools such as iTOL (Letunic and Bork, 2024), allowing for more tailored and focused analyses. Similarly, motif-related output files can be used with tools like WeLogo (Crooks et al., 2004) or Seq2Logo (Thomsen and Nielsen, 2012), enabling further customization and in-depth motif analysis. Additionally, intermediate files are preserved by default, allowing for inspection and troubleshooting at each stage of the analysis. Solutions for pairwise sequence comparison and visualization already exist (e.g., SDT (PMID: 25259891) and EFI-EST (PMID: 37356897)) and were not the focus of CRESSENT.

## 3. RESULTS

### 3.1 Rep and Cap Proteins Database

After acquiring ssDNA viral sequences from the ICTV database, we aimed to evaluate their similarity by constructing sequence similarity networks (SSNs) for both Rep and Cap proteins. For this, sequences from each dataset were concatenated and submitted to the EFI-EST (Oberg et al., 2023) platform for SSN generation, and the resulting networks were visualized using Cytoscape (Shannon et al., 2003). To construct SSNs specifically for Rep domains, sequences were first grouped by viral family and aligned using the align module from Cressent. Sequence logos were then generated via the WebLogo (Crooks et al., 2004) tool to pinpoint the position of the Walker A motif within each family. Based on the identified motif positions, regular expressions (regex) were created using a custom script. These regex patterns were subsequently used with the motif module to split each family’s sequences into functional domains (HUH and S3F domains). The extracted domain sequences were then concatenated and analyzed in EFI-EST to produce domain-specific SSNs, which were again visualized in Cytoscape. Networks at a 95% similarity threshold were examined, and associated metadata were used to annotate and color nodes by viral family, facilitating comparative analysis across groups (Fig. 2A-D).

**Fig 2:**
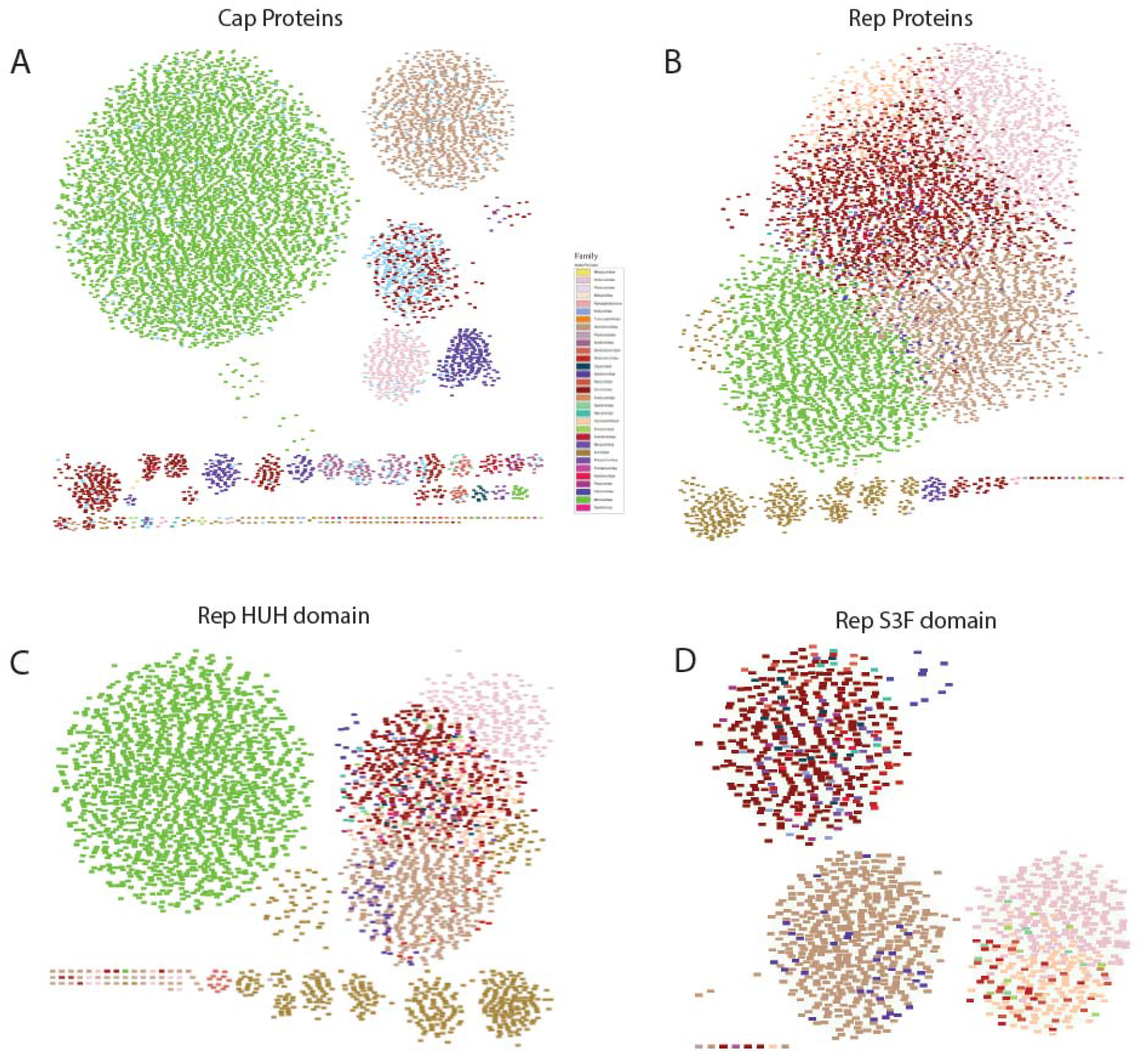
Protein sequence similarity network (SSN) of Cap and Rep (and domains) proteins. The SSN was generated using EFI-EST. Sequences sharing 95% identity are conflated as a single node and visualized by Cytoscape. Nodes are colored by family.

Cap protein sequences (Fig. 2A) exhibited more cohesive clustering overall compared to Rep proteins (Fig. 2B). However, when analyses focused on individual domains (HUH and S3F), Rep proteins formed clearer, more distinct clusters (Fig. 2C and 2D). Therefore, users should exercise caution when using full-length Rep sequences from diverse ssDNA virus families in a single analysis. We recommend conducting domain□level analysis and treating each Rep domain separately for accurate comparative studies.

### 3.2 CRESSENT can produce reproducible results

Sequences from two independent studies were employed to demonstrate the utility of CRESSENT in facilitating the annotation of Rep and Caps genes in two putative viral families, Naryaviridae (Zhang et al., 2025) and Genomoviridae (Leal Rodríguez et al., 2024). These viral sequences were identified and annotated using Cenote-Taker3 after undergoing quality trimming, assembly, and clustering. Figures 3 and 4 summarize the possible outputs produced by CRESSENT.

**Fig 3:**
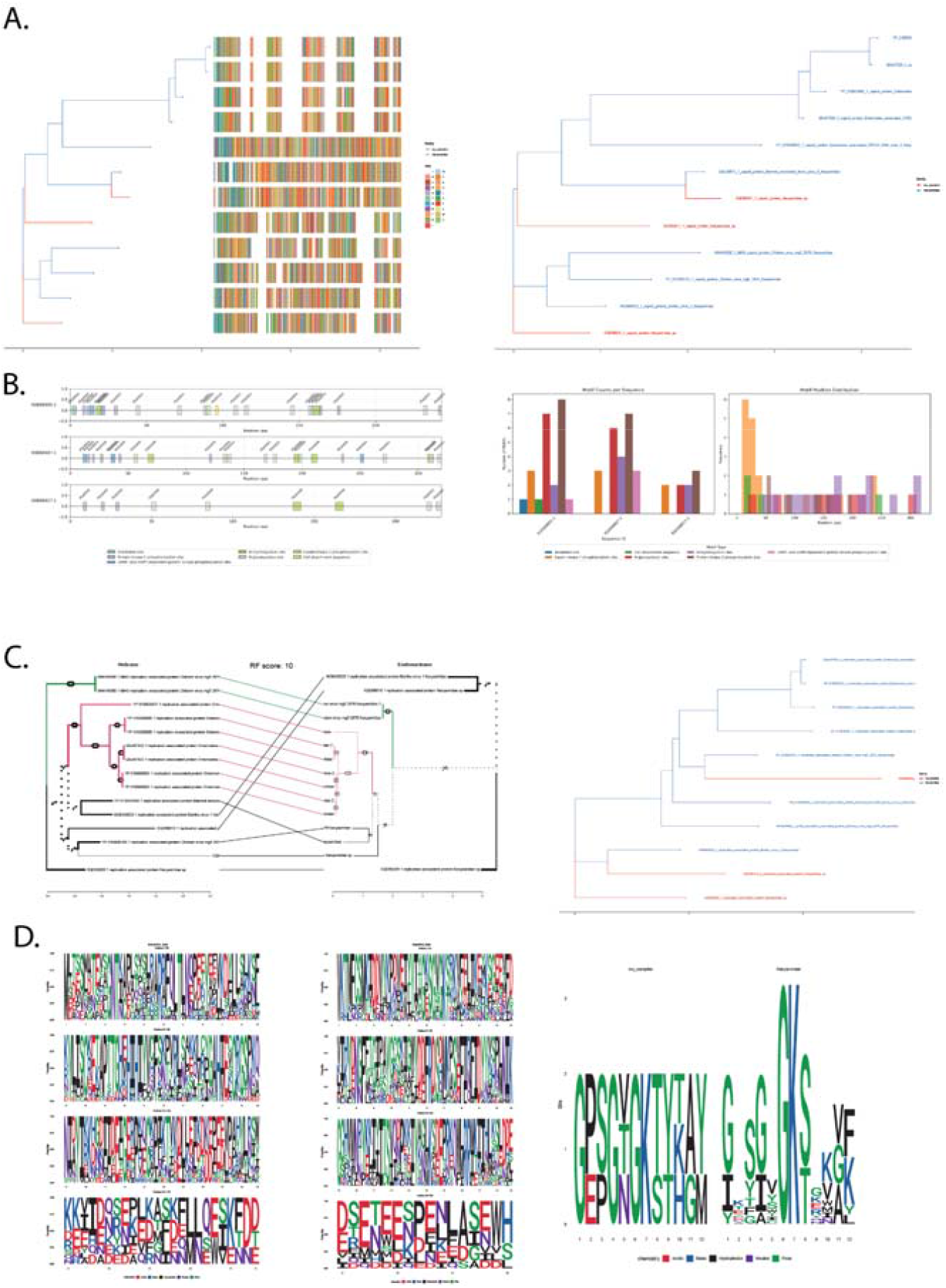
Visualizations produced by CRESSENT using Naryaviridae sequences. (A) Phylogenetic trees of Cap proteins (red = study samples; blue = custom DB sequences). (B) Gene visualization of the motifs found with the *de novo* motif module. (C) Tanglegram and phylogenetic tree of the Rep proteins. (D) Sequence logos of the HUH domain, the S3F domain, and the Walker A motif of Rep proteins.

**Fig 4:**
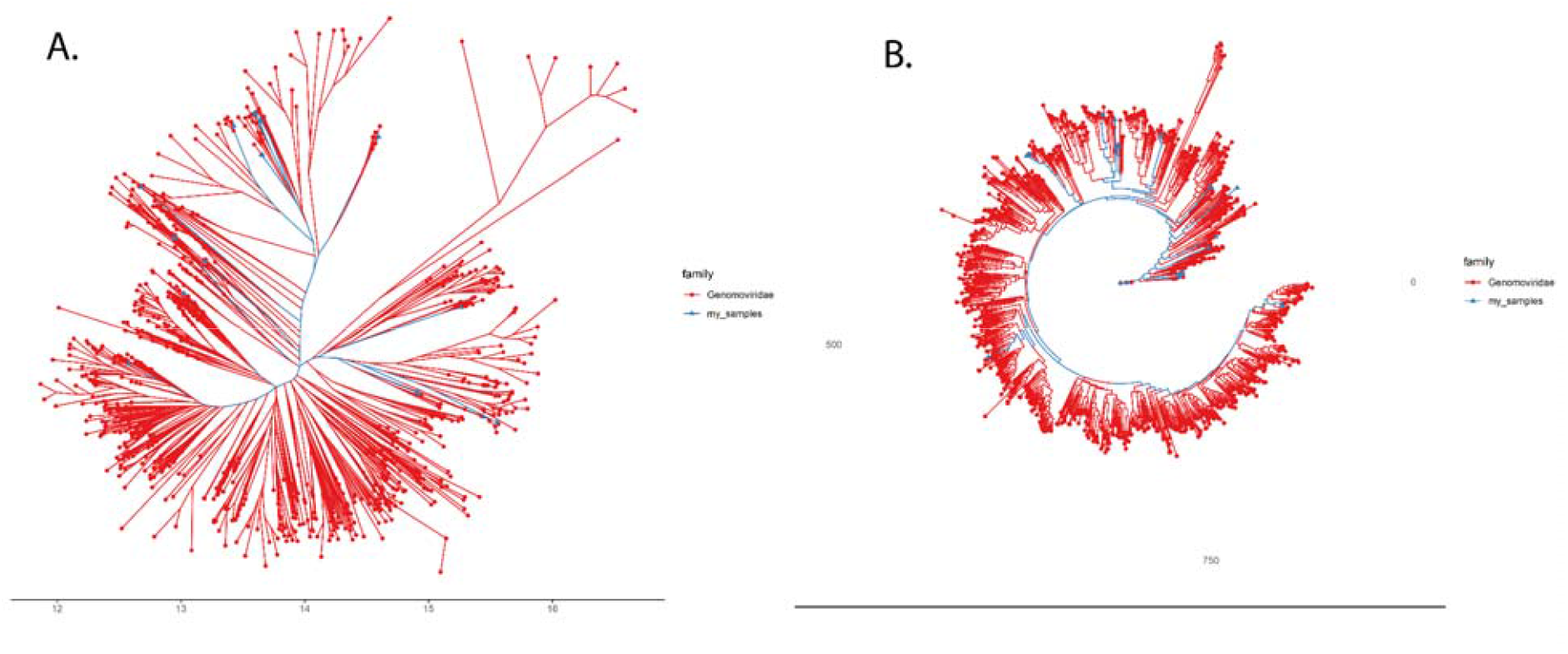
Unrooted (B) and circular (A) phylogenetic trees of the Genomoviridae Cap genes (blue = study samples; red = custom DB sequences).

## 4. CONCLUSION

The integration of multiple analysis methods provides researchers with a powerful toolkit for characterizing these important viral groups. The emphasis on data cleaning, visualization, and flexible workflow design enhances the utility of CRESSENT for diverse research applications. We anticipate that this tool will contribute to a more standardized and reproducible approach to ssDNA virus analysis, facilitating comparative studies and advancing our understanding of viral diversity and evolution. Importantly, the modular architecture of this tool was specifically designed to facilitate continuous improvement and expansion. This extensibility ensures that the tool can evolve alongside the rapidly advancing field of viral metagenomics, providing a sustainable platform for ssDNA virus analysis for years to come.

